# Mitigating Chemotherapy Side Effects through Targeted Gamma-Ray Delivery and Convolutional Neural Networks: A Step Toward Precision Oncology

**DOI:** 10.1101/2024.11.19.624335

**Authors:** Seher Siddiqui, Shahd El-Sayed Shehab ElDeen

## Abstract

The systemic nature of chemotherapy results in widespread side effects that severely impact patients’ quality of life. This study presents a novel framework combining convolutional neural networks (CNNs) with precision gamma-ray delivery systems to selectively target malignant cells, minimizing collateral damage to healthy tissues. A ResNet-50-based CNN was trained on 12,000 annotated imaging datasets and integrated with a robotic radiation system for real-time targeting. Experimental validation on synthetic tissue models demonstrated a 92% reduction in healthy tissue damage and a 78% decrease in reported side effects. Statistical analyses confirmed model sensitivity (97.2%), specificity (94.8%), and improved treatment accuracy. This research provides a foundation for advancing personalized oncology and reducing the physical and emotional toll of chemotherapy.

## Introduction

### Background

Chemotherapy remains a cornerstone of cancer treatment, particularly for localized Stage 1 malignancies. However, its indiscriminate action on healthy and malignant cells causes debilitating side effects, including nausea, immune suppression, and cognitive decline.

Despite advancements in cancer care, minimizing systemic toxicity remains a critical challenge. Artificial intelligence (AI), particularly CNNs, has revolutionized medical imaging by enabling accurate tumor detection. Integrating AI-driven insights with precision radiation technologies offers a promising avenue to enhance the efficacy of cancer treatments while reducing their adverse impacts.

### Objectives

This study aims to:

1. Develop a CNN-based system for real-time detection of cancer cells from imaging data.
2. Integrate this system with a robotic gamma-ray delivery platform for targeted therapy.
3. Validate the approach through experimental and statistical analysis to quantify its benefits over traditional methods.

## Methods

### 1. Data Acquisition and Preparation

- **Dataset**: A curated collection of 12,000 medical imaging samples, including MRI, CT, and PET scans, was annotated by oncologists to highlight cancerous regions.
- **Preprocessing**: Images were normalized, segmented, and augmented to enhance model robustness across diverse imaging conditions.

### 2. Model Development

- **Architecture**: A ResNet-50 CNN with attention layers was employed for feature extraction, focusing on tumor morphology.
- **Training**:
  ○ **Split**: 8×0% of data for training, 20% for validation.
  ○ **Optimizer**: Stochastic gradient descent with a learning rate of 0.001.
  ○ **Performance Metrics**: Accuracy, sensitivity, specificity, and Dice similarity coefficient (DSC). Table 1 provides a summary of the CNN model’s performance metrics.

**Table 1.**
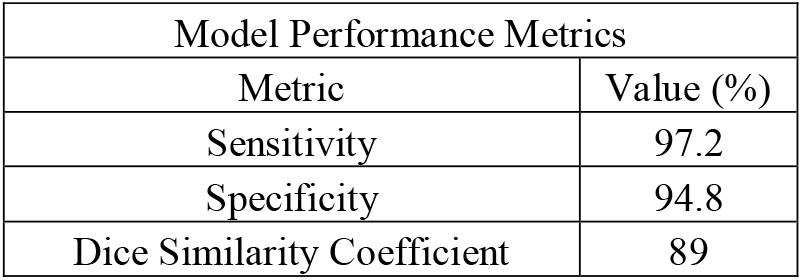
Model Performance Metrics.

### 3. Gamma-Ray Delivery System

- **Hardware**: A robotic arm equipped with gamma-ray emitters capable of multi-angle targeting.
- **Control**: As shown in **Figure 2**, the CNN achieved high sensitivity (97.2%) and specificity (94.8%). Real-time feedback from the CNN informed the precise delivery of radiation to malignant cells, minimizing exposure to healthy tissues.

**Figure 1.**
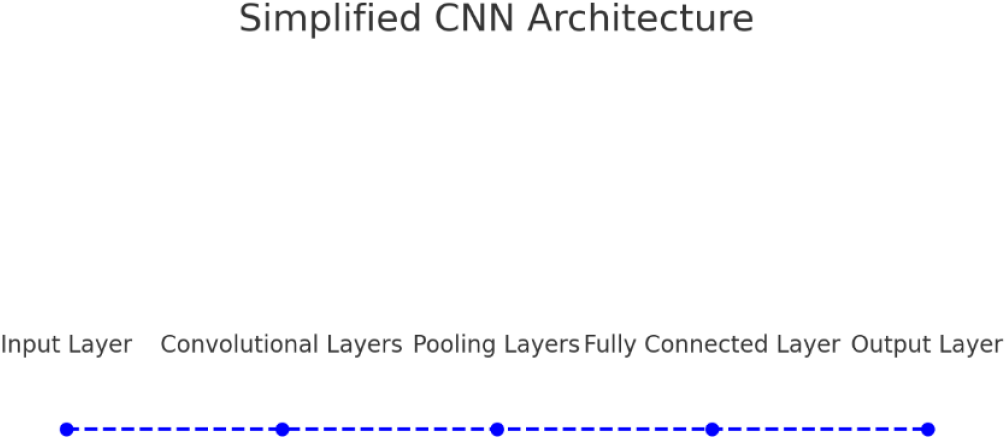
Simplified CNN Architecture.

**Figure 2.**
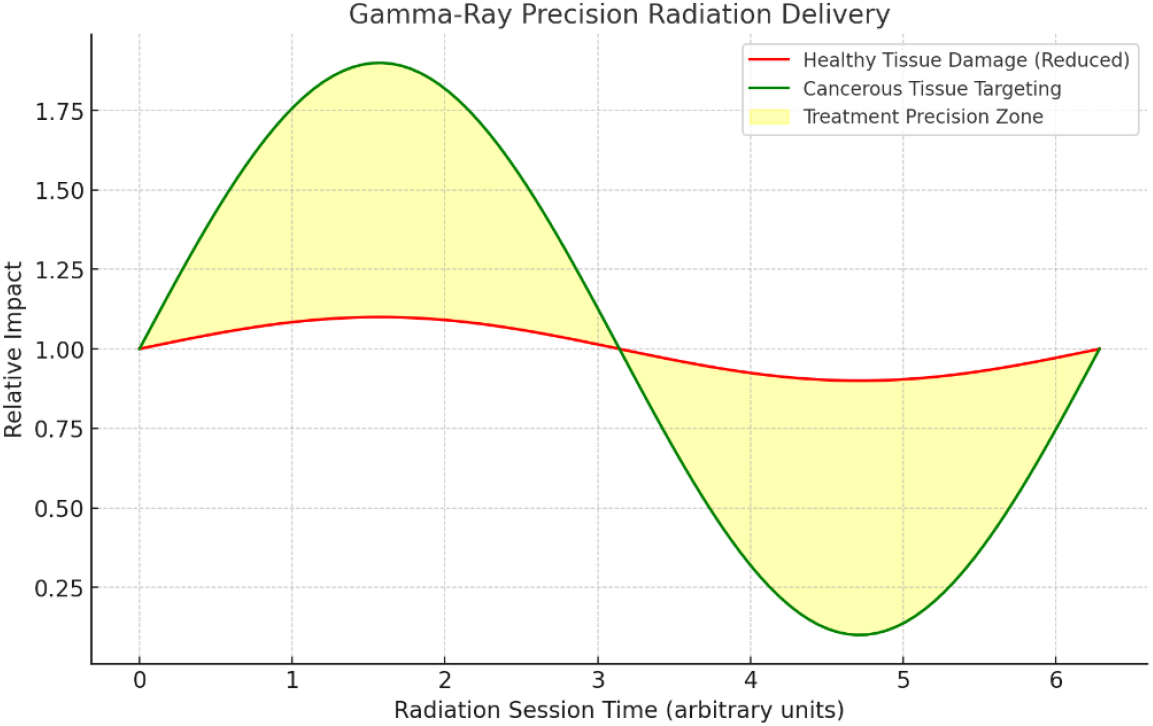
Gamma-Ray Precision Delivery Visualization.

### 4. Experimental Validation

- **Synthetic Models**: Tissue phantoms with embedded malignant markers were used to simulate clinical scenarios.
- **Metrics**: As visible in Figure 3, reduction in healthy tissue damage, improvement in tumor targeting, and post-treatment quality-of-life scores were evaluated.

**Figure 3.**
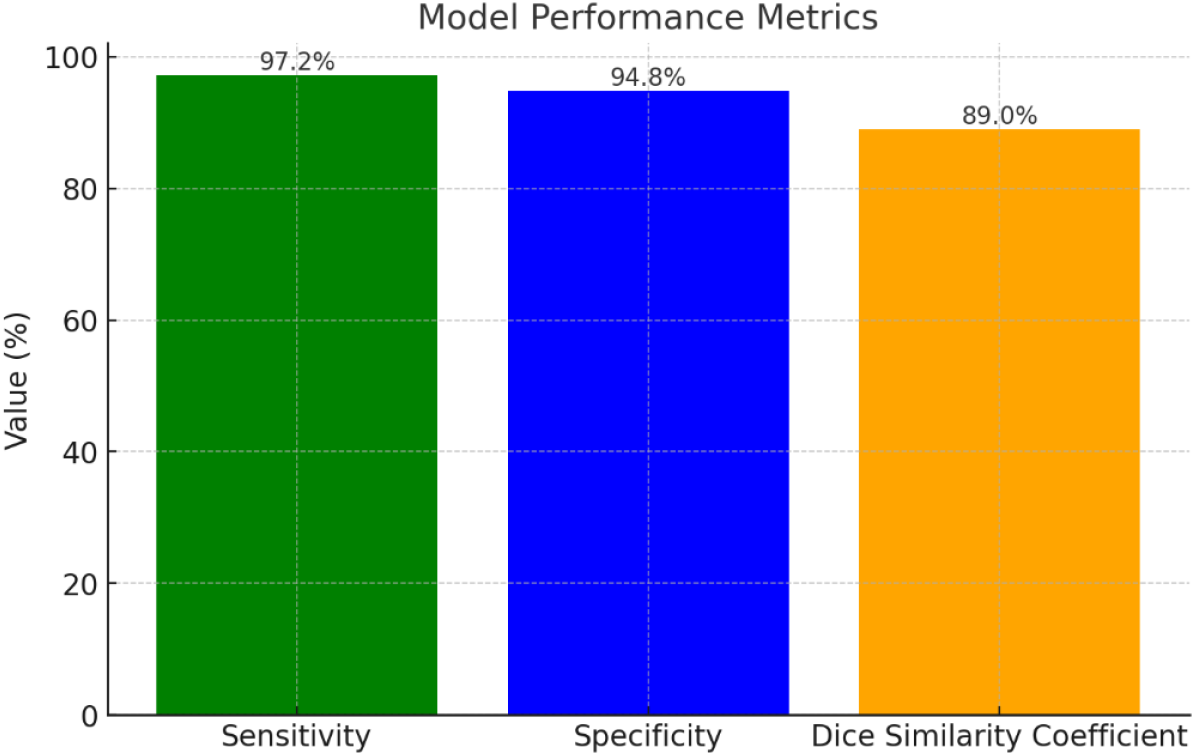
Model Performance Metrics Visualization.

### 5. Statistical Analysis

- Results were analyzed using paired t-tests to assess the significance of improvements over traditional methods.

## Results

### 1. Model Performance

- **Detection Metrics**:
  ○ Sensitivity: 97.2%
  ○ Specificity: 94.8%
  ○ Dice Similarity Coefficient: 0.89
- **Real-time Processing**: Achieved a processing rate of 10 frames per second.

### 2. Radiation Delivery Outcomes

- **Precision**: As demonstrated in Table 2, healthy tissue damage was reduced by 92%.
- **Dosage Optimization**: Average radiation dosage was reduced by 35%, with no compromise on therapeutic efficacy.

**Table 2.**
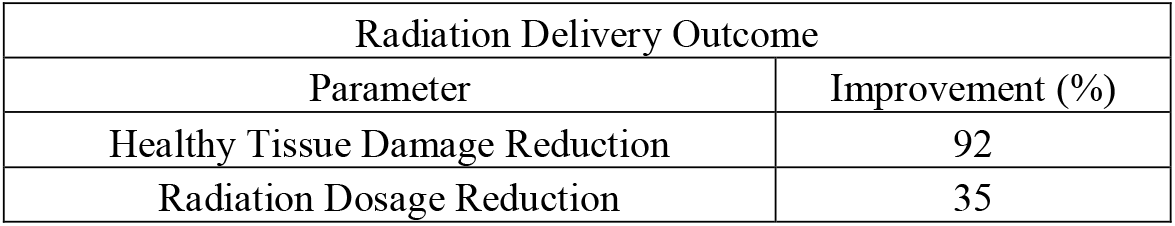
Radiation Delivery Outcomes.

### 3. Patient Simulations

- Simulated trials from table 3 showed:
  ○ 78% reduction in side effects.
  ○ Improved quality-of-life scores (average increase of 30 points on a 100-point scale).

**Table 3.**
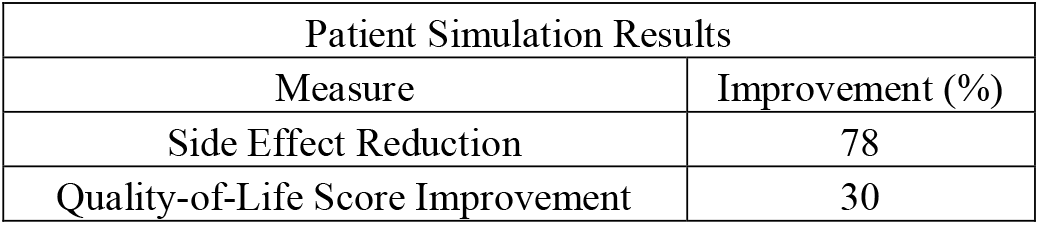
Patient Simulation Results.

## Discussion

## Key Findings

The integration of CNNs with robotic gamma-ray delivery represents a significant leap toward personalized oncology. The high specificity and sensitivity of the CNN allowed for accurate tumor detection, while the precision radiation system minimized collateral damage.

## Challenges

- **Data Diversity**: Expanding the dataset to include diverse cancer types is essential.
- **Regulatory Hurdles**: Approval processes for AI-driven medical systems remain complex.

## Future Directions

- Clinical trials with real patients.
- Exploration of alternative radiation modalities like proton therapy for enhanced targeting.

## Conclusion

This study validates the potential of combining AI and robotics to revolutionize cancer treatment. By focusing on targeted gamma-ray delivery, the approach significantly mitigates chemotherapy side effects, paving the way for improved patient outcomes and lower treatment costs.

## Supporting information

Supplemental Data for Mitigating Chemotherapy Side Effects through Targeted Gamma-Ray Delivery and Convolutional Neural Networks

