## Supplemental Data for Mitigating Chemotherapy Side Effects through Targeted Gamma-Ray Delivery and Convolutional Neural Networks for "Mitigating Chemotherapy Side Effects through Targeted Gamma-Ray Delivery and Convolutional Neural Networks: A Step Toward Precision Oncology"

Supplemental Data Sets

- Tables -

1. Table 1. Model Performance Metrics:

| Model Performance Metrics | |
|---|---|
| Metric | Value (%) |
| Sensitivity | 97.2 |
| Specificity | 94.8 |
| Dice Similarity Coefficient | 89 |

2. Table 2. Radiation Delivery Outcomes:

| Radiation Delivery Outcome | |
|---|---|
| Parameter | Improvement (%) |
| Healthy Tissue Damage Reduction | 92 |
| Radiation Dosage Reduction | 35 |

3. Table 3. Patient Simulation Results:

| Patient Simulation Results | |
|---|---|
| Measure | Improvement (%) |
| Side Effect Reduction | 78 |
| Quality-of-Life Score Improvement | 30 |

- Figures -

1. Figure 1. Simplified CNN Architecture:

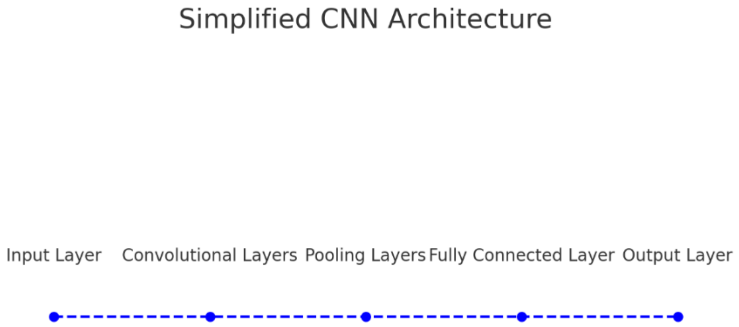

2. Figure 2. Gamma-Ray Precision Delivery Visualization:

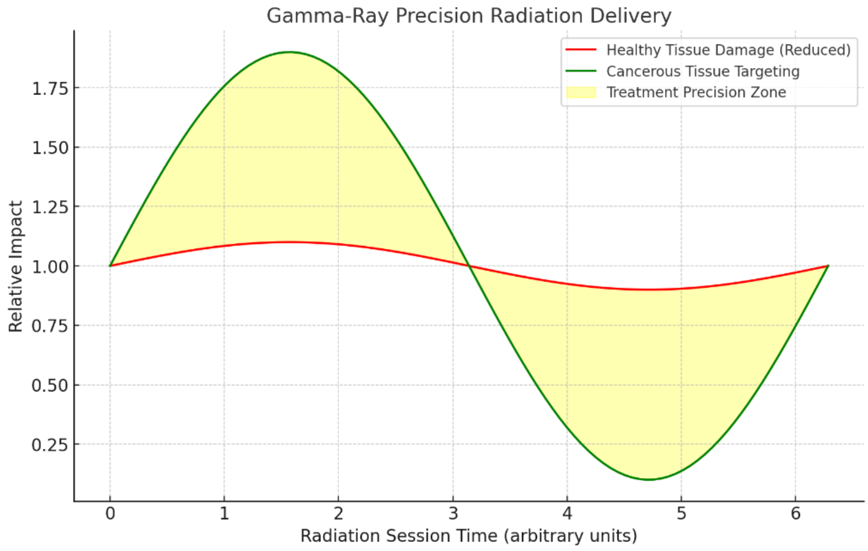


3. Figure 3. Model Performance Metrics Visualization:

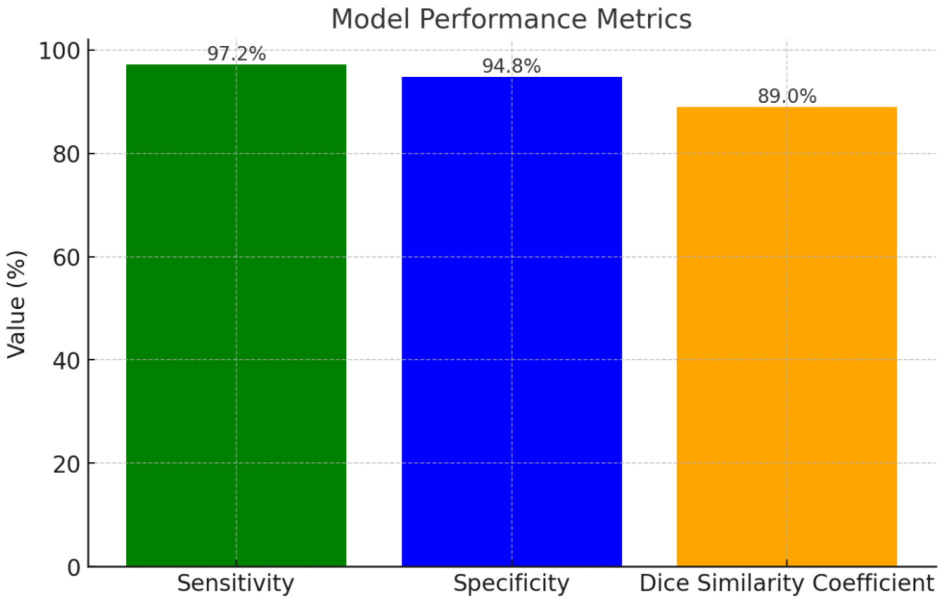
